# The social and spatial ecology of dengue presence and burden during an outbreak in Guayaquil, Ecuador, 2012

**DOI:** 10.1101/112185

**Authors:** Catherine A. Lippi, Anna M. Stewart-Ibarra, Ángel G. Muñoz, Mercy J. Borbor, Raúl Mejía, Keytia Rivero, Katty Castillo, Washington B. Cárdenas, Sadie J. Ryan

## Abstract

Dengue fever, a mosquito-borne viral disease, is an ongoing public health problem in Ecuador and throughout the tropics, yet we have a limited understanding of the disease transmission dynamics in these regions. The objective of this study was to characterize the spatial dynamics and social-ecological risk factors associated with a recent dengue outbreak in Guayaquil, Ecuador. We examined georeferenced dengue cases (n = 4,248) and block-level census data variables to identify potential social-ecological variables associated with the presence and burden of dengue fever in Guayaquil in 2012. We applied LISA and Moran’s I tests to analyze hotspots of dengue cases and used multimodel selection in R computing language to identify covariates associated with dengue incidence at the census zone level. Significant hotspots of dengue transmission were found near the North Central and Southern portions of Guayaquil. Significant risk factors for presence of dengue included poor housing conditions (e.g., poor condition of ceiling, floors, and walls), access to paved roads, and receipt of remittances. Counterintuitive positive correlations with dengue presence were observed with several municipal services such as garbage collection and access to piped water. Risk factors for the increased burden of dengue included poor housing conditions, garbage collection, receipt of remittances, and sharing a property with more than one household. Social factors such as education and household demographics were negatively correlated with increased dengue burden. Our findings elucidate underlying differences with dengue presence and burden and indicate the potential to develop dengue vulnerability and risk maps to inform disease prevention and control - information that is also relevant for emerging epidemics of chikungunya and zika.

**Highlights:**

- In 2012, Guayaquil, Ecuador had a large outbreak of dengue cases
- Dengue case presence and burden exhibited spatial heterogeneity at the census block level
- Social-ecological drivers of case presence and burden differed in this outbreak, highlighting the need to model both types of epidemiological data
- Access to municipal resources such as garbage collection and piped water had counterintuitive relationships with dengue presence, but poor housing, garbage collection and remittances correlated to dengue burden.
- Our findings inform risk mapping and vector control and surveillance allocation, relevant to this and other concurrent emergent epidemics such as chikungunya and zika

## 1. Introduction

The public health sector in Latin America is facing the alarming situation of concurrent epidemics of dengue fever, chikungunya, and zika, febrile viral diseases transmitted by the mosquito vectors *Aedes aegypti* and *Aedes albopictus* (Dick et al., 2012; Muñoz, Thomson, Goddard, & Aldighieri, 2016; Pan American Health Organization & World Health Organization, 2016; Zambrano et al., 2016). Traditional surveillance and vector control efforts have been unable to halt these epidemics (Stewart Ibarra et al., 2014). Macro-level social-ecological factors have facilitated the global spread, co-evolution and persistence of the dengue viruses and vectors, including growing urban populations, global movement, climate variability, insecticide resistance, and resource-limited vector control programs (Stewart Ibarra et al., 2014). It is necessary to study effects of these drivers at the local level to understand the complex dynamics and drivers of disease transmission, which vary from region to region, thus allowing decision makers to more effectively intervene, predict, and respond to disease outbreaks (Stewart Ibarra et al., 2014).

Spatial epidemiological risk maps provide important information to target focal vector control efforts in high-risk areas, potentially increasing the effectiveness of public health interventions (Castillo, Körbl, Stewart, Gonzalez, & Ponce, 2011; Castillo, Katty C., 2011). Typically, historical epidemiological records are digitized to understand the spatial distribution of the burden of disease and the presence/absence of disease. Layers of social-ecological predictors (e.g., land use maps, socioeconomic census data) are incorporated to test the hypothesis that one or more predictors are associated with the presence /absence or burden of disease. This information allows decision makers to identify geographic areas to focus interventions (e.g., hotspots) and risk factors to target in integrated vector-control interventions (e.g., community health interventions for specific vulnerable populations). Spatial risk maps can also be integrated into disease early warning systems (EWS) to generate disease risk forecasts, such as seasonal risk maps. (Banaitiene, 2012; Kuhn, Katrin, Campbell-Lendrum, Diarmid, Haines, Andy, & Cox, Jonathan, 2005; Thomson, García Herrera, & Beniston, 2008; “WHO | Global Strategy for dengue prevention and control, 2012–2020,” n.d.). Previous studies indicate that associations between social-ecological risk factors and dengue transmission may vary by location and time, highlighting the need for local analyses of dengue risk (Almeida, Medronho, & Valencia, 2009; Castillo et al., 2011; de Mattos Almeida, Caiaffa, Assunção, & Proietti, 2007; Mondini et al., 2009; Stewart Ibarra et al., 2013; Stewart-Ibarra et al., 2014; Teixeira et al., 2002).

Since 2000, DENV 1-4 have co-circulated in Ecuador, presenting the greatest burden of disease in the lowland tropical coastal region (Alava, A., Mosquera, C., Vargas, W., & Real, J., 2005). Guayaquil, Ecuador, the focus of this study, is the largest city in the country and the historical epicenter of dengue transmission in the country. The first cases of autochtonous chikungunya cases were reported in Ecuador at the end of 2014, resulting in a major epidemic in 2015, with over 33,000 cases reported. The first cases of zika were confirmed in Ecuador on January 7, 2016, and to date (25 Aug 2016) 2,076 suspected cases of zika have been reported by the Ministry of Health (Pan American Health Organization & World Health Organization, 2016). Climate is an important source of predictability for these diseases, mainly because both the viruses and the vectors are sensitive to temperature. For example, anomalously high surface temperatures produced by a combination of natural climate variability and climate change have been suggested to have played a role in the present zika epidemic (Muñoz et al., 2016).

The aim of this study was to characterize the spatial dynamics of cases and demographic risk factors associated with a recent dengue epidemic (2012) in the coastal city of Guayaquil, Ecuador. This study builds on prior studies in Machala, Ecuador, that demonstrated the role of social determinants in predicting dengue risk at the household and city-levels, and contributes to a broader effort to strengthen surveillance capacities in the region in partnership with the Ministry of Health and the National Institute of Meteorology and Hydrology (INAMHI) (Stewart Ibarra et al., 2013; Stewart-Ibarra et al., 2014). This study is intended to both provide the much needed local-level social-ecological context, and to demonstrate the differences arising in inference from presence and burden of cases in these analyses.

## 2. Material and methods

In this study, we build on methodologies used in prior work in Machala, Ecuador (Stewart Ibarra et al., 2013; Stewart-Ibarra et al., 2014), using many of the same methods, for comparison, but adding an examination of burden, beyond just presence of dengue. Guayaquil is far larger than Machala (ca. 10 times the population size), and therefore an interesting contrast to examine in the same framework as the previous work.

### 2.1 Study Area

Dengue fever is hyper-endemic in Guayaquil, Guayas Province, a tropical coastal port city (pop. 2,350,915) (INEC, 2010), as well as the largest city in Ecuador (Fig. 1). There is a pronounced seasonal peak in dengue transmission from February to May, which follows the onset of the hot rainy season (Fig. 2). In Guayaquil, in 2012, more than 4,000 cases of dengue fever (and 79 DHF) were reported, marking the biggest dengue outbreak in recent years (Real, J. & Mosquera, C., 2012). There were 4,248 clinically reported cases of dengue fever - an incidence of 18.07 dengue cases per 10,000 population per year compared to an average incidence of 4.99 dengue cases per 10,000 population per year from 2000 to 2011 (Fig. 3) [22].

**Figure 1.**
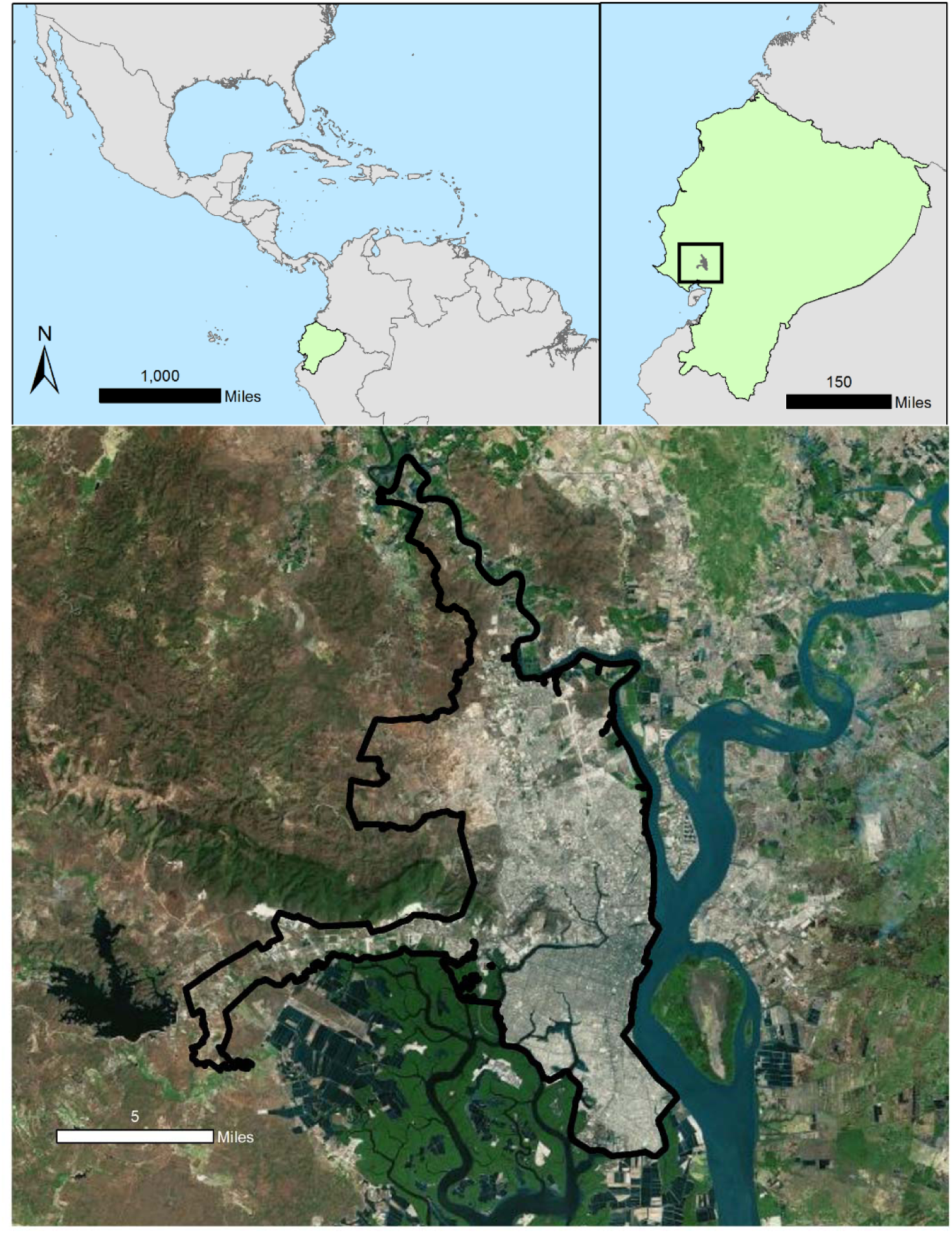
The study site, Guayaquil, is located within Guayas province in coastal Ecuador. Panel A shows the location of Ecuador (medium green) in S. America (bright green); Panel B shows the location of Guayaquil (orange) within Guayas Province (warm yellow), in Ecuador; Panel C shows the city of Guayaquil, detailing the census block structural layout, and showing where major and minor waterways exist surrounding the city limits. This figure was created in ArcGIS version 10.3.1 (*ArcGIS 10.3.1*, 2016) using shapefiles from the GADM database of Global Administrative Areas, version 2.8, freely available at gadm.org (“Global Administrative Areas (2015).,” 2015). Inland rivers and water bodies data are derived from the Digital Chart of the World (DCW)(US Military, Department of Defense (all branches), 1992). Census block outlines were digitized by INAMHI (National Institute of Meteorology and Hydrology) authors during the course of this project, as reported in the methods.

**Figure 2.**
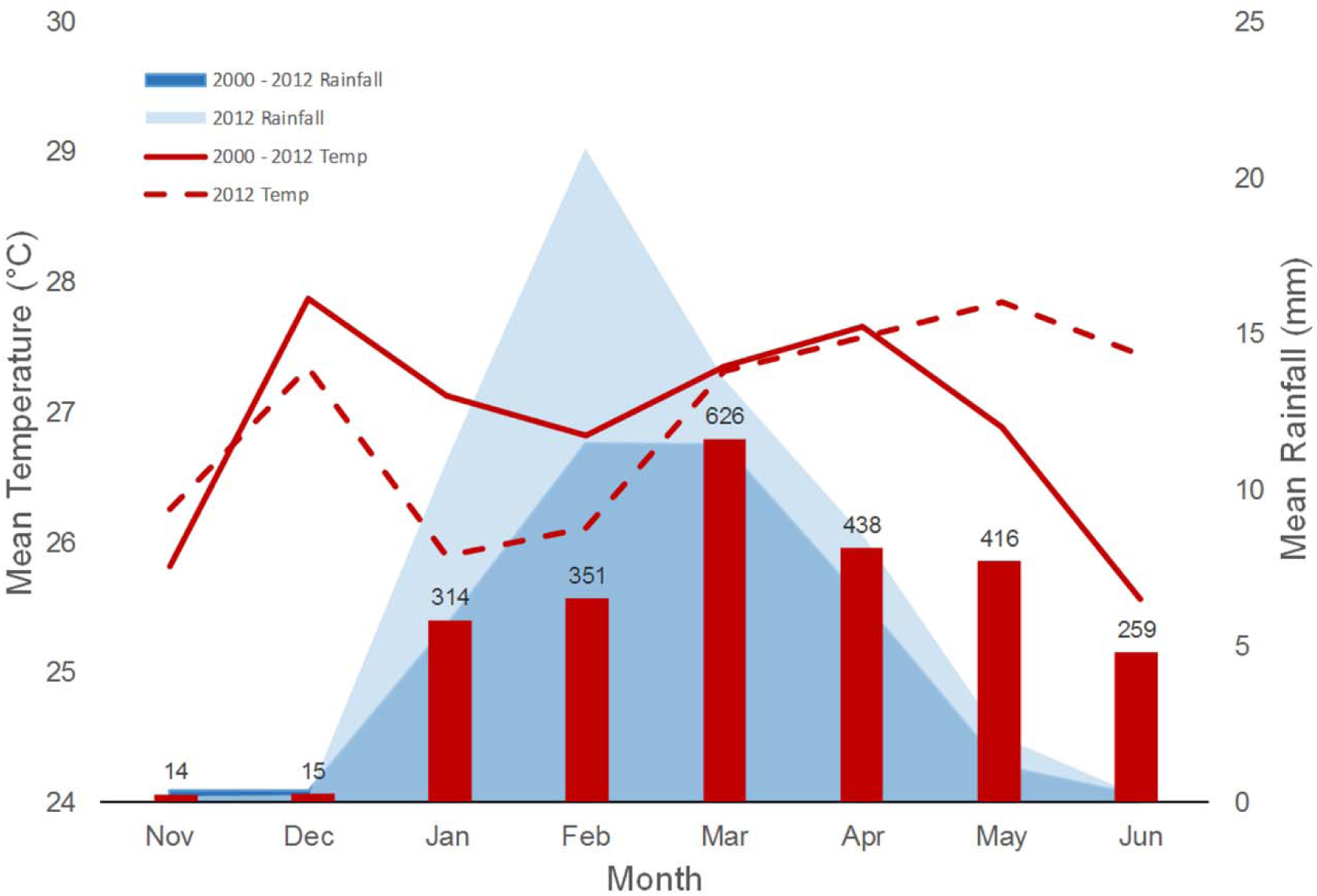
Climate trends in Guayaquil, Ecuador. The bold line shows the average temperature and rainfall from 2000 – 2012 in comparison with monthly average temperature for 2012 (dashed line). The red bars show monthly totals of confirmed dengue cases from 2012.

**Figure 3.**
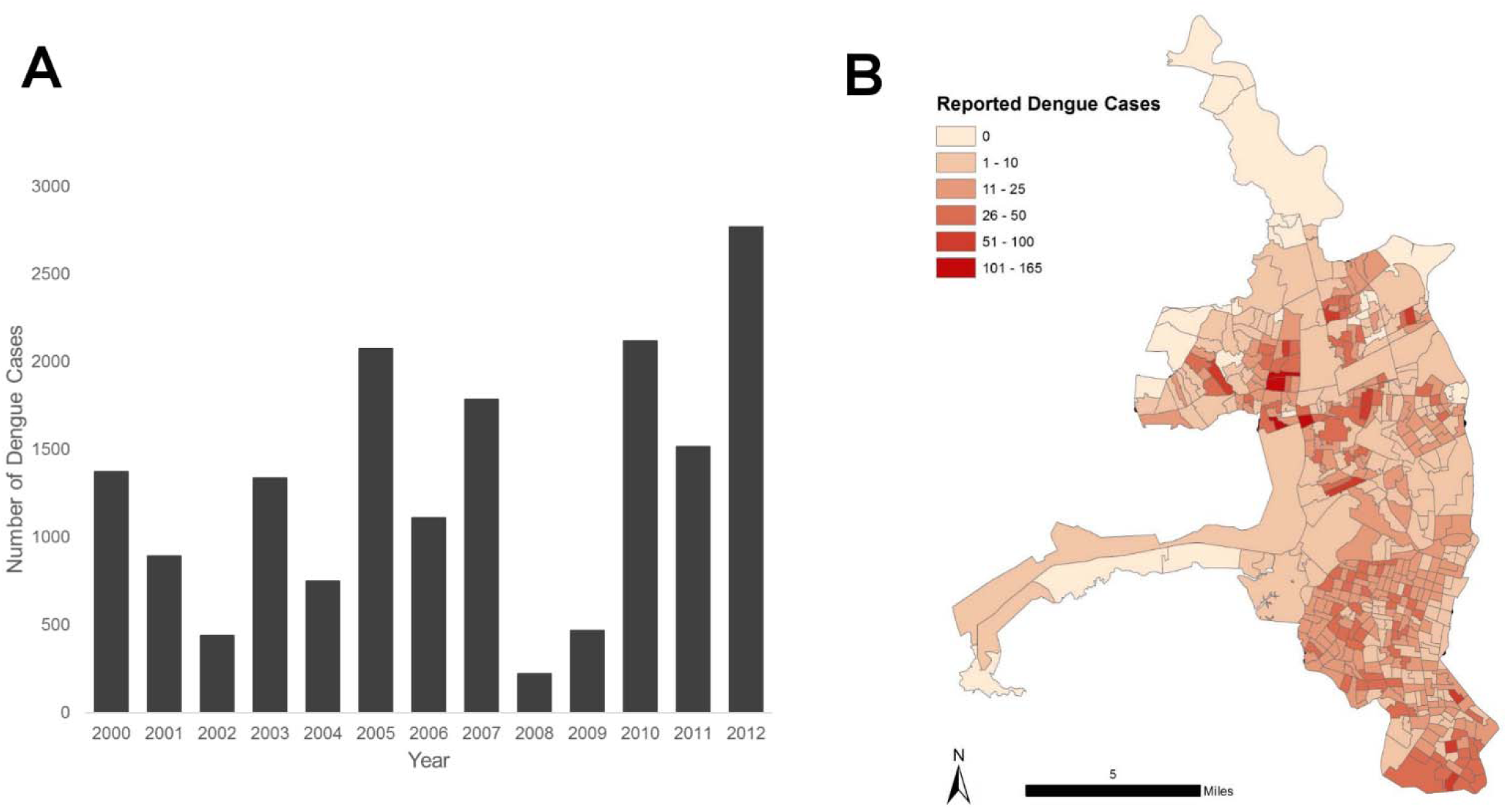
Total number of clinically diagnosed cases of dengue fever in Guayaquil, Ecuador (2000 – 2012) (A). Cases during the 2012 outbreak were reported throughout the city’s census zones (B). This figure was created in ArcGIS version 10.3.1 (*ArcGIS 10.3.1*, 2016) using shapefiles from the GADM database of Global Administrative Areas, version 2.8, freely available at gadm.org (“Global Administrative Areas (2015).,” 2015). Census block outlines were digitized by INAMHI (National Institute of Meteorology and Hydrology) authors during the course of this project, as reported in the methods.

### 2.2 Data Sources

Epidemiological data (dengue case reports for 2012) and national census data (2010) were examined to identify potential social-ecological variables associated with the presence and burden of dengue fever during the 2012 outbreak in Guayaquil, Ecuador. These data were provided by INAMHI through a collaborative project with the Ministry of Health, sponsored by the Ecuadorian government (Muñoz, Á.G., Stewart-Ibarra, A.M., & Ruiz-Carrascal, D., 2013). No formal ethical review was required as the data used in this analysis were de-identified and aggregated to the census zone level, as described in the following.

#### 2.2.1 Epidemiological data

For the analyses presented here, INAMHI provided de-identified georeferenced dengue cases from Guayaquil in 2012 (n = 4,248), aggregated and mapped to census zone polygons to protect the identity of individuals (Hartter, Ryan, MacKenzie, Parker, & Strasser, 2013) (Supplemental Data File 1). This map was generated from individual records of clinically suspected and confirmed cases of dengue fever and DHF (aggregated as total dengue fever) reported to a mandatory disease surveillance system operated by the Ministry of Health, and included 15.03% of total dengue cases confirmed by the Ministry of Health in Ecuador for 2012 (n= 16,544) (Ministerio de Salud Pública, 2013). Dengue cases included in this study were defined based on clinical diagnosis rather than laboratory confirmed cases due to the low rate of laboratory confirmation.

#### 2.2.2 Social-ecological risk factors

We extracted individual and household-level data from the 2010 Ecuadorian National Census (INEC, 2010) to assess social-ecological variables associated with the presence and burden of dengue (Table 1). We selected variables that have been previously described and used in similar epidemiological studies (Stewart-Ibarra et al., 2014). We created a normalized housing condition index (0 to 1, where 1 is the worst) by combining three housing variables: the condition of the roof, condition of the walls, and condition of the floors. We recoded selected census variables and calculated parameters as the percent of households or percent of the population per census zone (n = 484).

**Table 1.** Socio-ecological parameters (mean and standard deviation - SD) tested in logistic regression and negative binomial model searches to respectively predict presence of dengue and severity of outbreak.

| Parameter | Mean | SD |
| --- | --- | --- |
| **Housing conditions** |  |  |
| House condition index (HCI),<br>0 to 1, where 1 is poor condition | 0.27 | 0.12 |
| More than four people per bedroom | 16.78 % | 0.08 |
| People per household | 3.88 | 0.34 |
| Municipal garbage collection | 93.06 % | 0.15 |
| People in household drink tap water | 76.85 % | 0.09 |
| Piped water inside home | 77.03 % | 0.31 |
| Municipal sewage | 62.52 % | 0.39 |
| Access to paved roads | 80.06 % | 0.25 |
| More than one household per structure | 1.90 % | 0.01 |
| Unoccupied households | 16.08 % | 0.56 |
| Rental homes | 1.55% | 0.17 |
| **Demographics** |  |  |
| Receive remittances | 8.85 % | 0.04 |
| People emigrate for work | 1.88 % | 0.01 |
| Mean age of the head of the household (years) | 45.69 | 4.54 |
| Mean household age (years) | 29.36 | 4.29 |
| Proportion of household under 15 years of age | 28.34 % | 0.06 |
| Proportion of household under 5 years of age | 9.31 % | 0.03 |
| Head of the household has primary education or less | 30.94 % | 0.15 |
| Head of household has secondary education | 31.73 % | 0.07 |
| Head of household has post-secondary education | 25.77 % | 0.21 |
| Afro-Ecuadorian | 10.13 % | 0.07 |
| Head of the household is unemployed | 26.84 % | 0.06 |
| Head of the household is a woman | 33.29 % | 0.04 |

#### 2.2.3 Climate data

INAMHI provided rainfall and 2-meter temperature station data at monthly scale for the period 1981-2012. The long-term means were computed for both variables, and monthly values for the year 2012 were compared with those climatological values (Fig. 2). A complementary analysis was performed to understand the behavior of these two variables during 2012, using sea-surface temperature fields from both the Pacific and the Atlantic Oceans (ERSST version 4, (Huang et al., 2015), and vertically integrated moisture fluxes computed using the NCEP-NCAR Reanalysis Project version 2 (Kanamitsu et al., 2002).

### 2.3 Statistical Analyses

#### 2.3.1 Spatial analyses

We applied Moran’s I with inverse distance weighting in ArcGIS (ver. 10.3.1) (*ArcGIS 10.3.1*, 2016) to disease rates derived from epidemiological dengue case and population census data to test the hypothesis that dengue cases were non-randomly distributed in space. Moran’s I is a global measure of spatial autocorrelation, that provides an index of dispersion from -1 to +1, where -1 is dispersed, 0 is random, and +1 is clustered. We identified the locations of significant dengue hot and cold spots using Anselin’s Local Moran’s I with inverse distance weighting, in ArcGIS (*ArcGIS 10.3.1*, 2016). The Local Moran’s I is a local measure of spatial association (LISA) (Anselin, 2010) used to identify significant clusters (hot or cold spots) and outliers (e.g., nonrandom groups of neighborhoods with above or below the expected dengue prevalence). Previous studies have used global Moran’s I and LISA statistics to test the spatial distribution of dengue transmission (Oliveira, Ribeiro, & Castillo-Salgado, 2013), including in Ecuador (Castillo et al., 2011), allowing for comparison between studies.

#### 2.3.2 Social-ecological risk factors

We analyzed social-ecological variables associated with the presence and burden of dengue fever, including population density, human demographic characteristics, and housing condition (Table 1), were explored. We hypothesized that the presence or absence of dengue fever cases and the severity of the outbreak were associated with one or more of these social-ecological factors. Each factor comprised a suite of census variables, representing testable variable ensemble hypotheses in a model selection framework.

Two model searches were performed in R, using ‘glmulti’ for multimodel selection (Calcagno & Mazancourt, 2010). The first search was to determine which census factors were influencing the presence or absence of dengue in Guayaquil, specifying a logistic modeling distribution in a Generalized Linear Model (GLM) framework (GLM, family=binomial, link=logit). The second model search examined which census factors were influencing outbreak severity, defined as the localized concentration of dengue, using dengue case counts per census zone, offset by local population, as the dependent variable (GLM, family=negative binomial). Model searches were run until convergence using glmulti’s genetic algorithm (GA) (Calcagno & Mazancourt, 2010). Models were ranked based on Akaike’s Information Criterion (AIC) corrected for small sample size (AICc). The top ranked model for each search was compared to its respective global model, which included all proposed variables as model parameters (Burnham, Anderson, & Burnham, 2002). Parameter estimates and 95% confidence intervals (CI) were calculated for variables in the top ranked model from each search. Variance inflation factors (VIF) were calculated to assess multi-collinearity and model dispersion.

## 3. Results

### 3.1 Spatial analyses

Dengue incidence in census zones ranged from 0 cases (n = 88 zones) to 160 cases per 10,000 population (n = 1 zone) (Fig. 4). Dengue cases were significantly clustered (Moran’s I = 0.066, p < 0.05), and the LISA analysis indicated significant dengue hotspots (n = 30 high-high census zones) in the North Central and Southern areas of the city, in addition to a small number of significant outliers (n = 3 high-low neighborhood, n = 7 low-high neighborhoods) (p < 0.05, Fig. 4).

**Figure 4.**
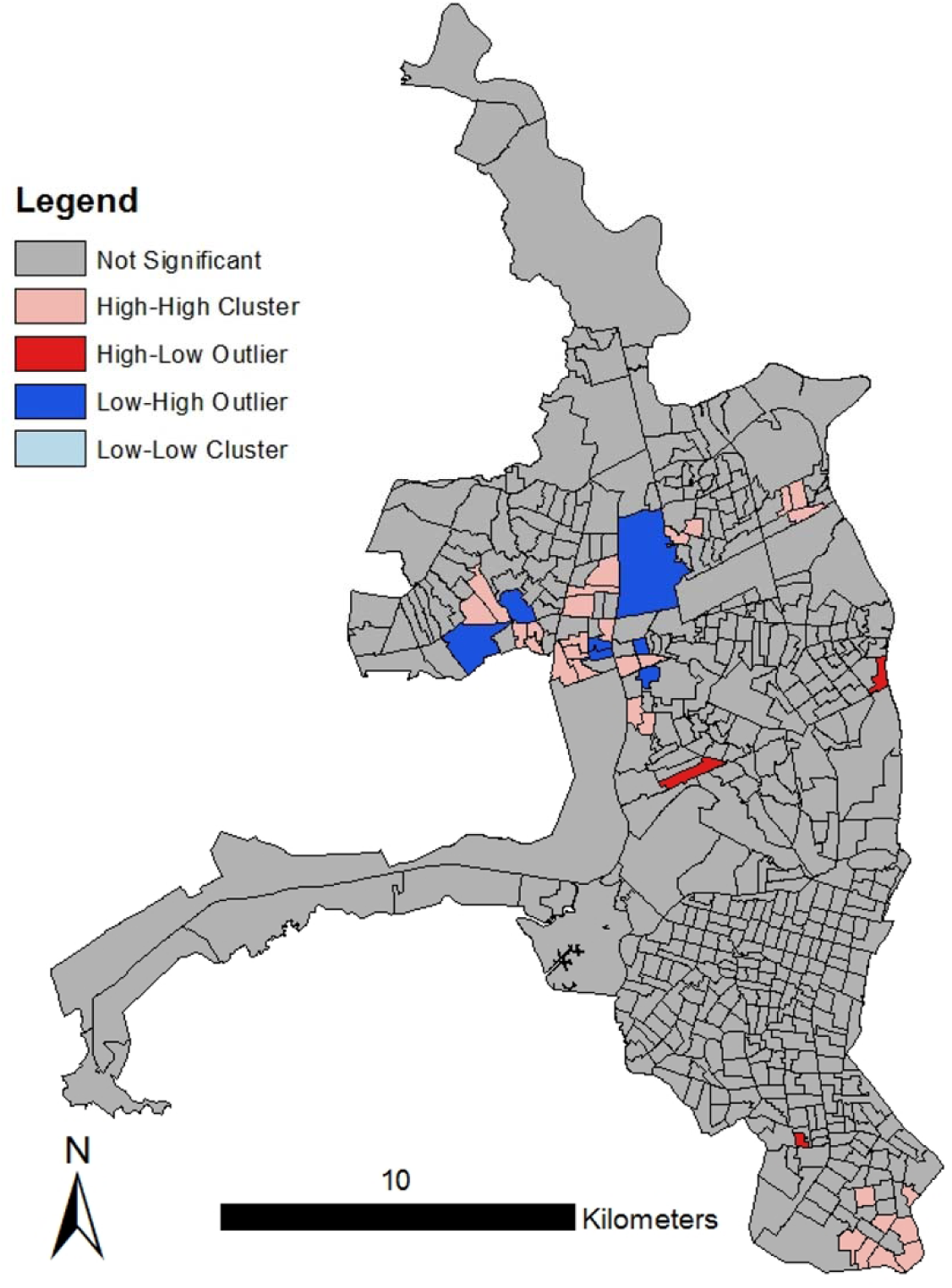
LISA analysis for the 2012 Guayaquil outbreak. Cases of dengue were significantly clustered in the North Central and Southern areas of the city. This figure was created in ArcGIS version 10.3.1 (*ArcGIS 10.3.1*, 2016) using shapefiles from the GADM database of Global Administrative Areas, version 2.8, freely available at gadm.org (“Global Administrative Areas (2015).,” 2015). Census block outlines were digitized by INAMHI (National Institute of Meteorology and Hydrology) authors during the course of this project, as reported in the methods.

### 3.2 Social-ecological risk factors

The most important risk factors associated with the presence of cases of dengue fever were poor housing conditions (e.g., poor structural condition of the floor, roof, and walls) and the proportion of households that received remittances. Other significant risk factors positively associated with the presence of dengue included greater access to municipal services (sewerage, piped water, garbage collection), fewer households that drink tap water, and lower proportion of Afro-Ecuadorians in the local population (Table 2). Ten additional models were found to be within 2 AICc units of the top model (Supplemental Table 1).

**Table 2.**
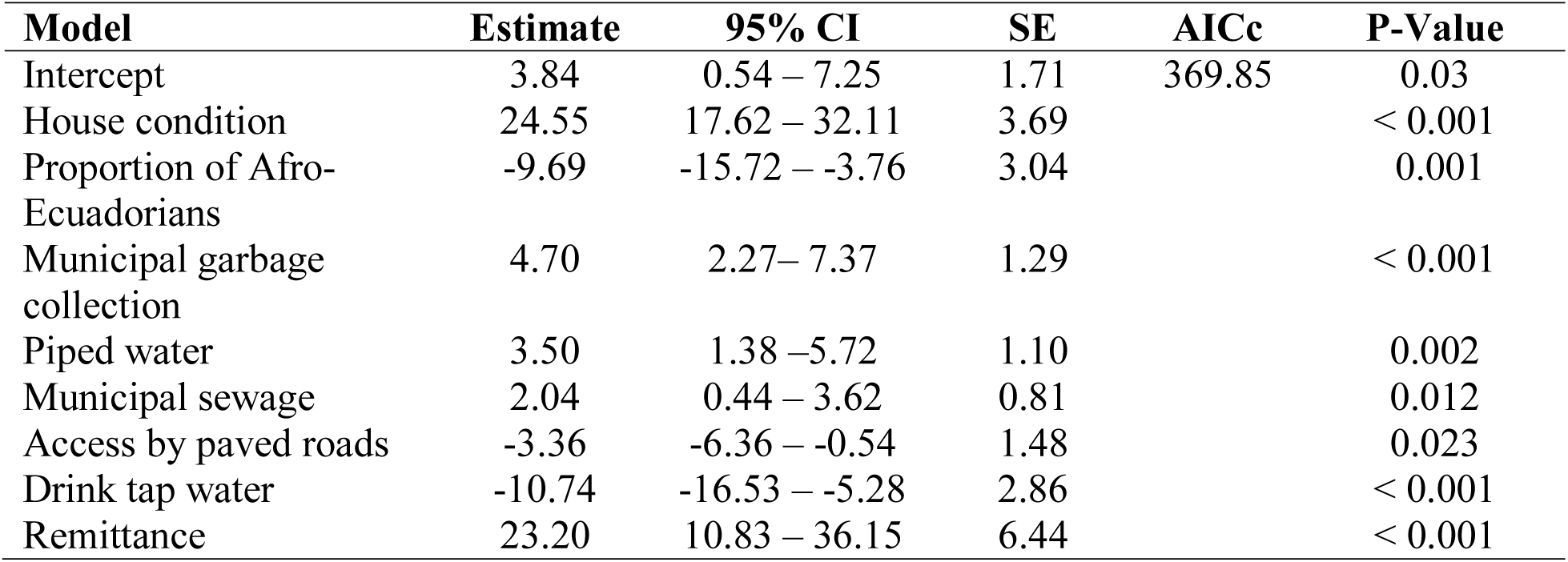
Top logistic regression model used in determining which social-ecological factors are important to dengue presence.

Poor housing condition was also the most important risk factor associated with the severity of localized dengue outbreaks in Guayaquil. Other factors positively associated with the number of dengue cases per census zone included lower proportion of heads of household with postsecondary and primary education, lower proportion of Afro-Ecuadorians in the population, lower proportion of household members under 15 years of age, older age of the heads of household, greater access to municipal garbage collection, a greater proportion of housing structures with more than one household, and a greater proportion of families receiving remittances (Table 3). Twenty-nine additional models were found to be within 2 AICc units of the top model (Supplemental Table 2).

**Table 3.**
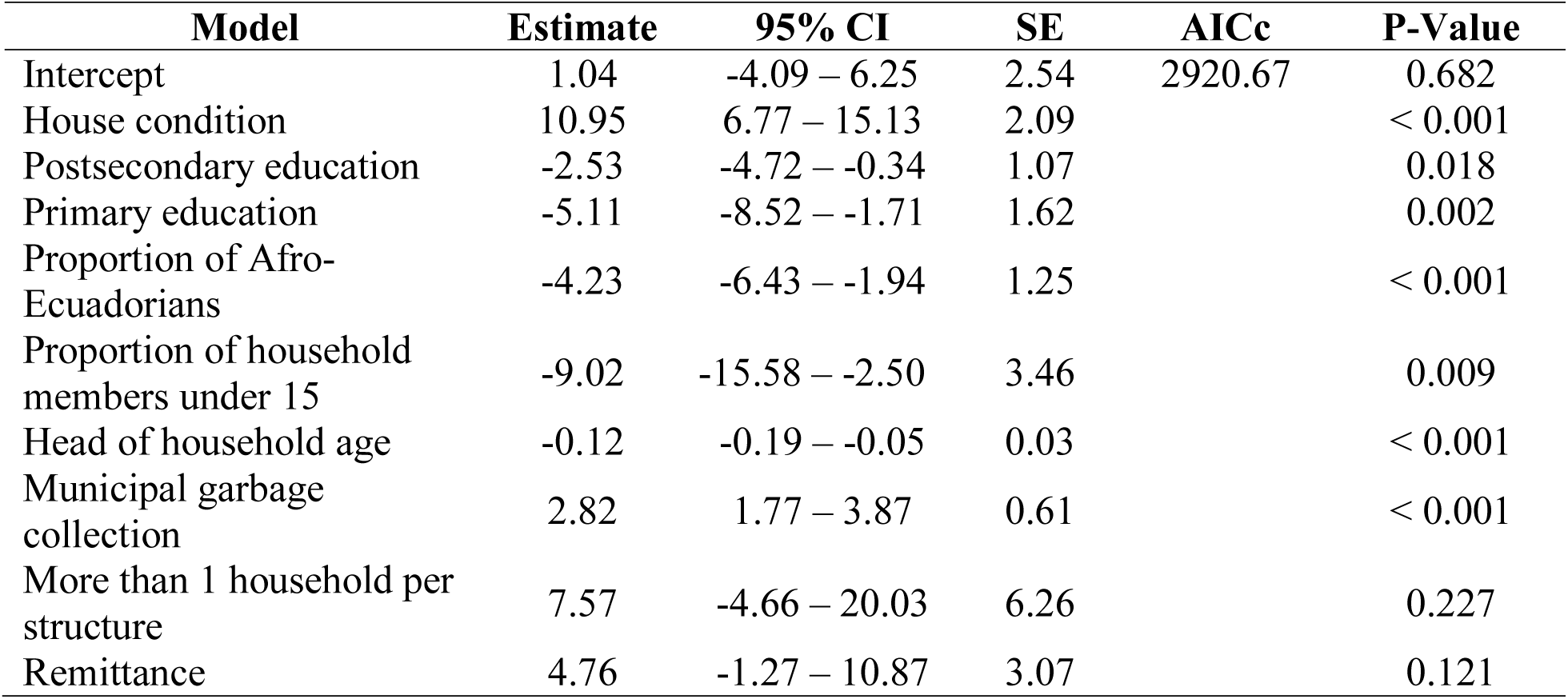
Top negative binomial model used in determining which social-ecological factors are important to dengue burden.

Results from the VIF analysis showed that 17 of the 23 tested variables had VIF scores under 10, indicating a fair degree of collinearity among certain predictors within census categories (e.g. measures of education and household age structure showed some correlation) Collinear variables were included in the multimodel searches as the main concern with inflated VIF scores is large error terms, not the coefficient estimates. Collinear variable suites were shown to be significant in many top models even with conservative model search criteria in place.

### 3.3 Climate analysis

The 2012 outbreak occurred toward the end of a weak La Niña event (2011/2012), with a peak of reported dengue cases around March, just after the precipitation peak of February brought anomalously high rainfall (approximately twice as much as the typical values for Guayaquil), and concurrent with an increase in temperatures from below-normal to normal seasonal values (Fig. 2). Although identified as a weak La Niña due to the behavior of the sea-surface temperature anomalies in the Equatorial Pacific (see Supplemental Figure 1, “climate” a,c,d), anomalously high moisture fluxes continuously arrived to coastal Ecuador from the Pacific during January and March (see Supplemental Figure 1, “climate” b,d,e), providing suitable conditions for the above-normal rainfall amounts observed during the season in Guayaquil.

## 4. Discussion

Since the 1980s, febrile illnesses transmitted by *Aedes aegypti* and *Aedes albopictus* (dengue fever, chikungunya, zika fever) have been increasing in incidence and distribution despite ongoing vector control interventions (Dick et al., 2012; San Martín et al., 2010; “WHO | Global Strategy for dengue prevention and control, 2012–2020,” n.d.). Targeted interventions and new surveillance strategies are urgently needed to halt the spread of these diseases. The results of this study highlight the need to differentiate between disease burden and presence when developing risk maps, providing an important contribution to our understanding of the spatial dynamics of dengue transmission. This study also provides an important local-level characterization of transmission dynamics, which are complicated by the non-stationary relationships among apparent dengue infection, climate, vector, and virus strain dynamics (Cazelles, Chavez, McMichael, & Hales, 2005; Cummings et al., 2004; Reiner et al., 2014); and the geographic and temporal variation in the social-ecological conditions that influence risk (Almeida et al., 2009; de Mattos Almeida et al., 2007; Mondini et al., 2009; Teixeira et al., 2002).

### 4.1 Spatial characteristics

During the 2012 outbreak, we identified hotspots of dengue fever transmission in the North Central and Southern areas of the city of Guayaquil, where land use is a mix of densely populated urban neighborhoods, industrial lots, and parks. Although they have access to basic services, previous studies suggest that communities in the urban periphery in coastal Ecuador have limited social organization and interaction with local authorities (Stewart Ibarra et al., 2014). Vector control in these areas consists of larvicidal products distributed by public health workers, but these products must be applied by individual households. Although there has been no formal evaluation of public mosquito abatement, health workers have indicated that homeowners do not use provided larvicides (M Borbor-Cordova, *pers comm*.). It should be noted that these census data do not capture the quality of the access to services, for example, the frequency of interruptions in the piped water supply or the frequency of garbage collection, which have a direct effect on mosquito breeding sites. Previous studies have also found significant clustering of dengue transmission in urban landscapes (Almeida et al., 2009; Galli & Chiaravalloti Neto, 2008; Hu, Clements, Williams, & Tong, 2011; Vazquez-Prokopec, Kitron, Montgomery, Horne, & Ritchie, 2010). In Guayaquil, previous work also identified neighborhood-level dengue hot and cold spots, which shifted over a 5-year period, pointing to the importance of continued spatial surveillance, and tracking potential risk factor shifts (Castillo et al., 2011; Castillo, Katty C., 2011). Fine scale spatial and temporal clustering of dengue has also been seen in Thailand (Endy et al., 2002; Yoon et al., 2012), and in Peru, where urban spatial transmission dynamics have been linked to human movement patterns within the urban environment (Stoddard et al., 2009; Vazquez-Prokopec et al., 2009). Given the reported flight range of the *Aedes aegypti* vector of approximately 250m (Castillo et al., 2011; Castillo, Katty C., 2011), we suggest that a combination of vector flight range, and intra-urban human movement, may lead to moderate hotspot patterns, while enabling broad scale spread of dengue across Guayaquil.

### 4.2 Social-ecological risk factors

Poor housing condition was the most important risk factor for dengue transmission in Guayaquil, influencing both the presence of dengue cases and the localized burden of the outbreak. Dengue was more likely to be present in a census zone when housing structures (i.e. roofs, walls, and floors) were in poor condition, access to paved roads was limited, and the proportion of houses receiving remittances was high. The risk factors for higher dengue burden were poor housing condition, proportion of houses receiving remittances, and the number of dwellings housing more than one family. These results suggest that accessibility of households to mosquitoes via structural deficiencies, as well as the overall socioeconomic status of neighborhoods, played a role in the 2012 outbreak (Fig. 5).Although the link between poverty and dengue transmission is not well characterized, the relationship between poor housing structure and arbovirus transmission has been well documented (Bradley et al., 2013; Hiscox et al., 2013; Mulligan, Dixon, Joanna Sinn, & Elliott, 2015; Birgit H. B. Van Benthem et al., 2005). Following the economic crisis in the late 1990s, many Ecuadorians immigrated to the U.S., Spain, and other countries in Europe for work, resulting in fragmented households and communities. The role of immigration in urban dengue control and prevention should be explored further (S. Bertoli, Fernandez-Huertas Moraga, & Ortega, 2011; Simone Bertoli & Marchetta, 2014; Jokisch & Pribilsky, 2002). When modeling the presence of dengue, all top models included access to core municipal services such as garbage collection, sewage, access to piped water, and number of houses drinking tap water as positive predictors of dengue cases (Table 2, Supplement 1). Municipal garbage collection was also positively correlated with dengue burden in all top models (Table 3, Supplement 2). Previous studies in smaller communities have observed positive correlations between lack of services and dengue transmission, as poor sanitation and water storing habits in urban areas are well-documented for providing habitat for larval *Aedes* mosquitoes (Stewart Ibarra et al., 2013; Stewart-Ibarra et al., 2014). Although municipal services are known to reduce the amount of larval mosquito habitat, there is some evidence to suggest that heavily urbanized areas, like Guayaquil, provide ample habitat regardless of service availability (Mulligan et al., 2015). Municipal services in Guayaquil are spatially heterogeneous, but in general services are more widely available in densely populated areas of the city (Fig. 6). However access to service does not necessarily serve as an indicator for quality or frequency of services. Several studies have identified the interaction between local *Aedes* production and human population density as a key factor in triggering dengue outbreak events (Almeida et al., 2009; Padmanabha, Durham, Correa, Diuk-Wasser, & Galvani, 2012; Rodrigues et al., 2015; Schmidt et al., 2011). The observed counterintuitive findings may indicate that although access to service should reduce the amount of available habitat for larval mosquitoes, human population density and quality of service may play a larger role in urban transmission of dengue. Intermittent or interrupted service may in fact exacerbate local conditions for mosquito breeding habitat, by increasing standing refuse piles, or prolonging duration of standing refuse, and in the case of water, increase problematic storage habits.

**Figure 5.**
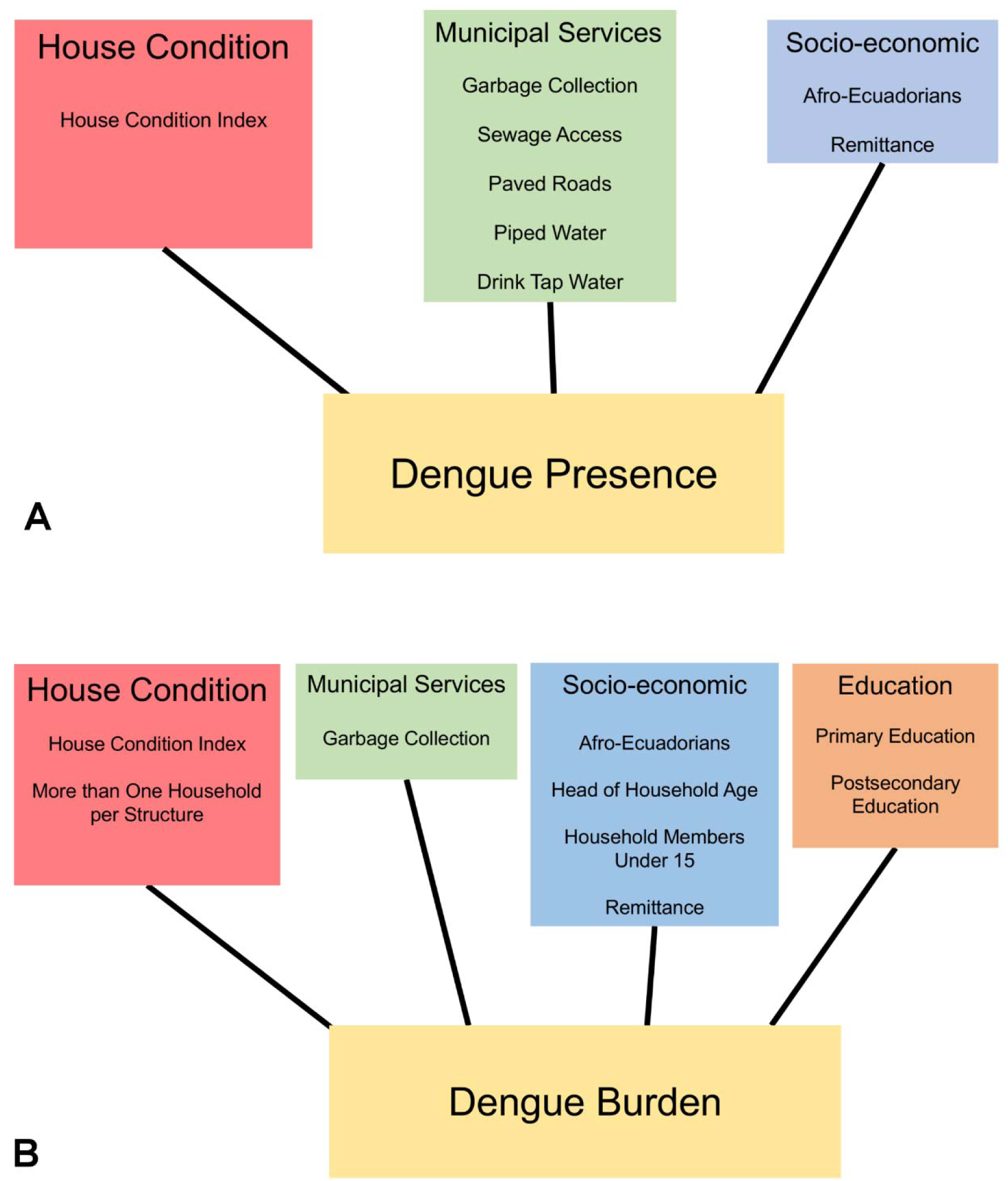
Conceptual diagrams highlighting the census variable suites that significantly affected dengue presence (A) and dengue burden (B) in Guayaquil, Ecuador during the 2012 outbreak.

**Figure 6.**
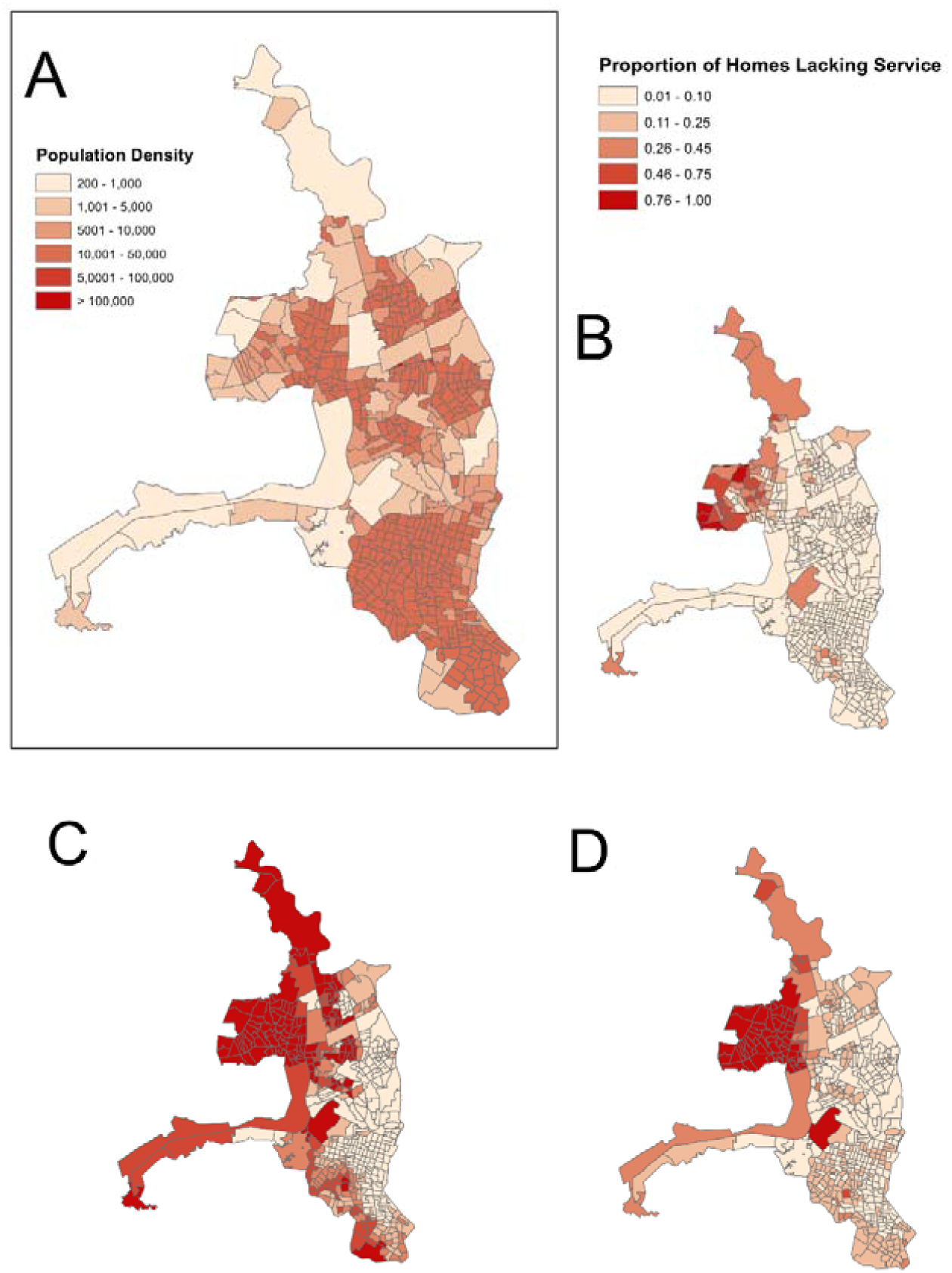
Population density (people per km^2^) of census zones in Guayaquil (A) shown against the proportion of homes lacking municipal garbage collection (B), lacking municipal sewage (C), and lacking piped water (D). Although dengue cases were reported in both densely and sparsely populated census zones, dengue hot spots were more associated with higher density zones (Fig.3), and the proportion of homes that lack basic municipal services tends to be higher in zones with lower population density. This may account for the counterintuitive model estimates associated with lack of these services (Tables 2 & 3). This figure was created in ArcGIS version 10.3.1 (*ArcGIS 10.3.1*, 2016) using shapefiles from the GADM database of Global Administrative Areas, version 2.8, freely available at gadm.org (“Global Administrative Areas (2015).,” 2015). Census block outlines were digitized by INAMHI (National Institute of Meteorology and Hydrology) authors during the course of this project, as reported in the methods.

Several demographic characteristics were found to be negatively correlated with dengue burden, i.e. age structure of households and access to primary and secondary education. Education, specifically knowledge about dengue, has been shown to influence the prevention practices of households and elimination of mosquito breeding sites (B. H. B. Van Benthem et al., 2002). Previous work in Machala, Ecuador, also revealed that household-level risk factors and perceptions of dengue risks vary with social and economic structures between communities (Stewart Ibarra et al., 2014). The proportion of Afro-Ecuadorians per census zone was associated with both lower dengue presence and burden, indicating the possibility of cultural and racial differences influencing localized transmission, or disproportionate case reporting.

Our findings on the social-ecological drivers of dengue fever transmission in Guayaquil are consistent with a prior study of household risk factors for the *Ae. aegypti* vector in the neighboring city of Machala, Ecuador. This previous study showed that housing condition and access to piped water inside the home were positively associated with the presence of *Ae. aegypti* pupae in the urban periphery of Machala (Stewart Ibarra et al., 2013). While the Machala study was in a smaller city, a different year, and different climate conditions, together these studies indicate the potential to target high risk households for vector control and dengue case management, using a locally adapted rapid survey of housing conditions.

The model selection framework used in this study is an effective strategy for exploratory studies to capture a large number of complex social-ecological processes. In contrast to traditional frequentist statistical approaches, a model selection approach enabled us to test multiple hypotheses simultaneously and identify potentially important variables for inclusion, not limited to significant variables determined by arbitrary *p*-values, or excluded due to collinearity before testing. Information theoretic or likelihood modeling approaches allow the modeler, who has *a priori* knowledge of the system, to make explicit informed decisions about which variables to include in testing in the model, and explore multiple compatible hypotheses rather than being limited to testing and excluding individual competing hypotheses. Additionally, the model search algorithms in the R package ‘glmulti’, facilitating exploration of all subsets of all possible models, is a more robust model selection procedure than stepwise regression techniques, which can lead to biased estimates (Calcagno & Mazancourt, 2010).

Guayaquil is a large, heterogeneous urban area, and there may be reporting bias of dengue cases especially in less populated areas with reduced access to medical care. However, reporting bias may not be as profound in Guayaquil as in other less-developed coastal cities in Ecuador. While the number of reported dengue cases was highest in densely populated census zones, cases were consistently reported throughout most of the city (Figure 3). Previous studies have shown that there is spatial and temporal variation in dengue hotspots within Guayaquil (Castillo et al., 2011; Castillo, Katty C., 2011). With multiple years of data at finer timescales, we could evaluate whether dengue transmission at the beginning of the dengue season or at the beginning of an epidemic is more likely to begin in neighborhoods with similar characteristics, to assess whether there are persistent high-risk, hotspot communities that trigger outbreaks. The analyses were limited by a lack of laboratory confirmation for cases, although efforts are ongoing to both improve dengue diagnostic infrastructure in the region, and to reduce the gap between epidemiological reporting and vector control interventions.

### 4.3 Climate conditions

Rainfall excess in 2012 produced moisture-saturated soils, formation of ponds of different sizes, water accumulation in a variety of containers, and other suitable conditions for vector proliferation. The transition to higher temperatures between February (rainfall maximum) and March is hypothesized here to have also contributed to the outbreak, as was the case for an analysis of similar conditions for a dengue outbreak in Machala [15]. We note that the fact that it was not a strong La Niña positively contributed to the occurrence of the dengue epidemic, as in this region, La Niña tends to be associated with a higher number of rainfall extreme events, leading to more runoff and a harder environment for mosquito breeding.

## 5. Conclusions

We found that census zones with certain social-ecological factors were more likely to report dengue cases during the 2012 outbreak in Guayaquil. We also demonstrated that the number and magnitude of variables influencing the presence versus the burden of dengue were fundamentally different, although some similarities were noted. The most important risk factors for both presence and increased burden of dengue during the outbreak included poor housing condition and the proportion of households receiving remittances. We also found evidence for spatial clustering of dengue fever transmission at the local sub-city level, highlighting areas for investigation into underlying causes of risk in these areas. Spatial clustering of dengue cases within Guayaquil and identification of social-ecological drivers for disease presence and burden indicate the potential to develop operational rapid survey tools to target high risk households, and the importance of spatially explicit analyses to inform surveillance and intervention efforts. This study contributes to ongoing efforts in developing early warning systems and predictive models for dengue in coastal Ecuador, undertaken by INAMHI and the Ministry of Health. These results help to inform the development of dengue vulnerability maps and data-driven dengue seasonal forecasts (Kuhn, Katrin et al., 2005).

## Abbreviations

AIC: Akaike’s Information Criterion
AICc: Akaike’s Information Criterion corrected for small sample size
CDC: Centers for Disease Control
CI: confidence intervals
DENV: dengue virus
DHF: dengue hemorrhagic fever
EWS: early warning systems
GA: genetic algorithm
GIS: geographic information system
GLM: Generalized Linear Model
INAMHI: National Institute of Meteorology and Hydrology
LISA: local indicators of spatial association
NCEP-NCAR: National Center for Environmental Protection – National Center for Atmospheric Research
VIF: variance inflation factors

## Acknowledgements

Many thanks to colleagues at the Ministry of Health and the National Institute of Meteorology and Hydrology for supporting ongoing climate–health initiatives in Ecuador. AGM used computational resources from the Observatorio Latinoamericano de Eventos Extraordinarios (www.ole2.org) and Centro de Modelado Científico (CMC), Universidad del Zulia.

## Declarations

### Ethics Approval

No formal ethical review was required as the data provided from INAMHI used in this analysis were de-identified and aggregated to the census zone level as described in the methodology.

### Data Availability

The data that support the findings of this study were made available through the Ecuadorian Ministry of Health and INAMHI, but restrictions apply to the availability of these data which were used in partnership for the current study. As such these data are not publicly available. Data are however available from the authors upon reasonable request and with permission of INAHMI.

### Authors’ contributions

AMSI, AGM, MJB, and RM conceived of the investigation. KR compiled the data used in analyses. AMSI, AGM, CAL, and SJR conducted analyses and drafted the manuscript. All co-authors assisted with interpretation of the data, providing feedback for this manuscript. All authors read and approved the final manuscript.

---

**Supplemental Figure 1.**
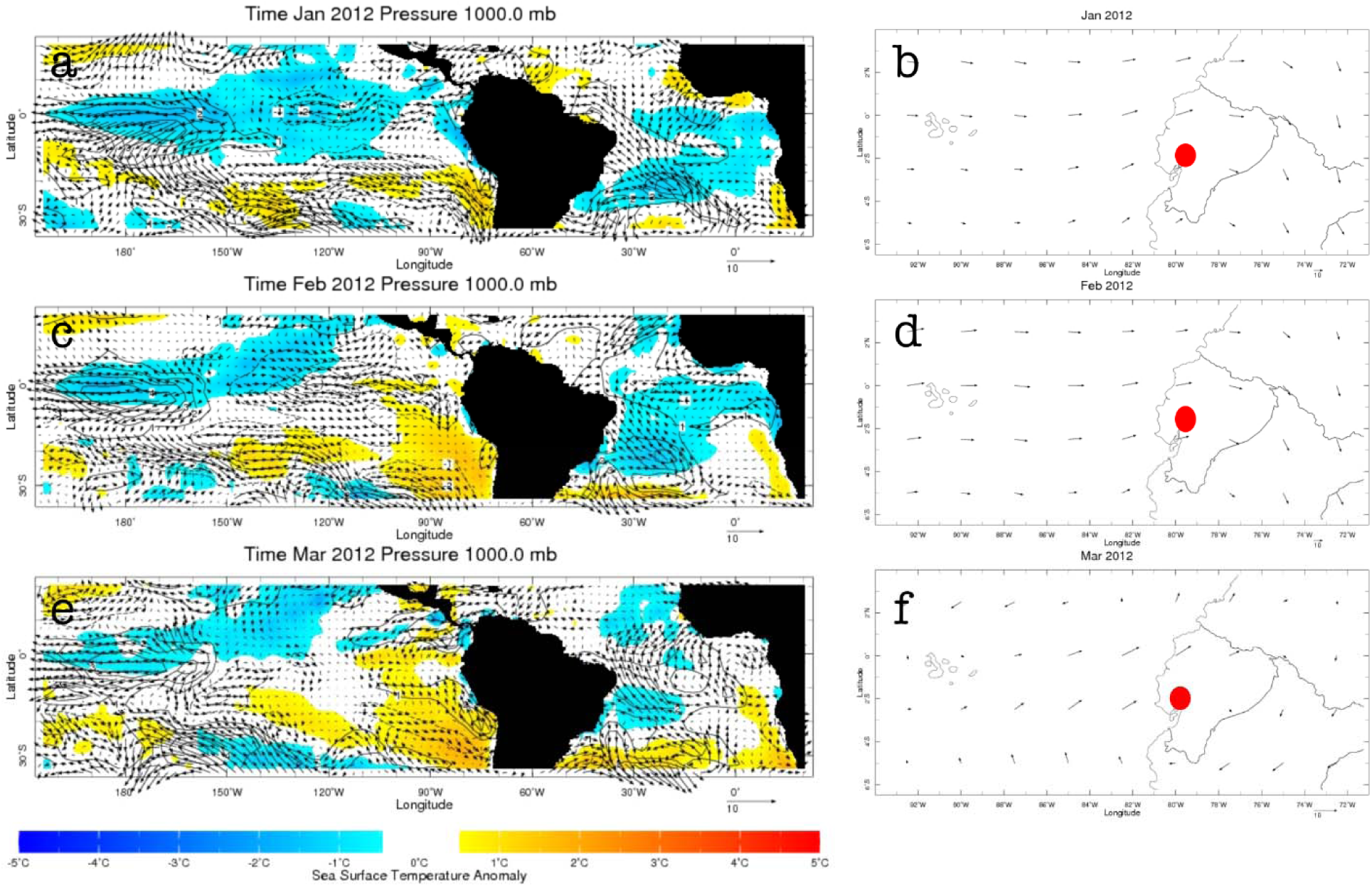
Sea-surface temperature anomalies (contours (°C)) and surface-level winds (vectors (m/s)) (panels a, c, e); vertically-integrated moisture flux anomalies (g/kg m/s) over Ecuador (panels b,d,f), during January, February and March 2012 (top, middle and bottom rows, respectively). Red dot indicates location of Guayaquil.

**Supplemental Table 1.** All logistic regression models for presence or absence of dengue cases ≤ 2 AICc units.

| Model | AICc | Weight |
| --- | --- | --- |
| x ~ 1 + afro + housecond + no_garbage + no_piped + no_sewage + no_pave + remit + drinktap | 369.85 | 0.11 |
| x ~ 1 + afro + unocc + housecond + no_garbage + no_piped + no_sewage + no_pave + remit + drinktap | 370.97 | 0.06 |
| x ~ 1 + afro + pplperhh + housecond + no_garbage + no_piped + no_sewage + no_pave + remit + drinktap | 371.07 | 0.06 |
| x ~ 1 + afro + fourpplbedrm + housecond + no_garbage + no_piped + no_sewage + no_pave + remit + drinktap | 371.10 | 0.06 |
| x ~ 1 + afro + housecond + no_garbage + no_piped + no_sewage + no_pave + gt1hh + remit + drinktap | 371.13 | 0.06 |
| x ~ 1 + afro + housecond + no_garbage + no_piped + no_sewage + no_pave + remit + drinktap + rental | 371.39 | 0.05 |
| x ~ 1 + prim + afro + housecond + no_garbage + no_piped + no_sewage + no_pave + remit + drinktap | 371.61 | 0.04 |
| x ~ 1 + afro + housecond + no_garbage + no_piped + no_sewage + no_pave + remit + emigr + drinktap | 371.68 | 0.04 |
| x ~ 1 + afro + womhead + housecond + no_garbage + no_piped + no_sewage + no_pave + remit + drinktap | 371.70 | 0.04 |
| x ~ 1 + afro + unemploy + housecond + no_garbage + no_piped + no_sewage + no_pave + remit + drinktap | 371.73 | 0.04 |
| x ~ 1 + sec + afro + housecond + no_garbage + no_piped + no_sewage + no_pave + remit + drinktap | 371.82 | 0.04 |

**Supplemental Table 2.** All negative binomial regression models for burden of dengue cases ≤ 2 AICc units.

| Model | AICc | Weight |
| --- | --- | --- |
| x ~ 1 + postsec + prim + afro + under15 + headhh_age + housecond + no_garbage + gt1hh + remit | 2920.67 | 0.03 |
| x ~ 1 + postsec + prim + afro + under15 + headhh_age + housecond + no_garbage + no_sewage + gt1hh + remit | 2920.71 | 0.03 |
| x ~ 1 + postsec + prim + afro + under15 + headhh_age + housecond + no_garbage + no_sewage + gt1hh + remit + drinktap + rental | 2920.76 | 0.03 |
| x ~ 1 + postsec + prim + afro + under15 + headhh_age + housecond + no_garbage + gt1hh + remit + drinktap + rental | 2921.05 | 0.03 |
| x ~ 1 + postsec + prim + afro + under15 + headhh_age + womhead + housecond + no_garbage + no_sewage + gt1hh + remit | 2921.16 | 0.03 |
| x ~ 1 + postsec + prim + afro + under15 + headhh_age + housecond + no_garbage + no_sewage + gt1hh + remit + rental | 2921.18 | 0.03 |
| x ~ 1 + postsec + prim + afro + under15 + headhh_age + womhead + housecond + no_garbage + gt1hh + remit | 2921.20 | 0.03 |
| x ~ 1 + postsec + prim + afro + under15 + headhh_age + fourpplbedrm + housecond + no_garbage + no_sewage + gt1hh + remit | 2921.20 | 0.03 |
| x ~ 1 + postsec + prim + afro + under15 + headhh_age + housecond + no_garbage + no_piped + no_sewage + gt1hh + remit + rental | 2921.48 | 0.02 |
| x ~ 1 + postsec + prim + afro + under15 + headhh_age + housecond + no_garbage + gt1hh + remit + drinktap | 2921.52 | 0.02 |
| x ~ 1 + postsec + prim + afro + under15 + hh_age + headhh_age + housecond + no_garbage + no_sewage + gt1hh + remit + drinktap + rental | 2921.60 | 0.02 |
| x ~ 1 + postsec + prim + afro + under15 + headhh_age + fourpplbedrm + housecond + no_garbage + gt1hh + remit | 2921.65 | 0.02 |
| x ~ 1 + postsec + prim + afro + under15 + hh_age + headhh_age + housecond + no_garbage + gt1hh + remit | 2921.66 | 0.02 |
| x ~ 1 + postsec + prim + afro + under15 + headhh_age + womhead + housecond + no_garbage + no_sewage + gt1hh + remit + drinktap + rental | 2921.77 | 0.02 |
| x ~ 1 + postsec + prim + afro + under15 + headhh_age + housecond + no_garbage + gt1hh + remit + rental | 2921.97 | 0.02 |
| x ~ 1 + postsec + prim + afro + under15 + headhh_age + fourpplbedrm + housecond + no_garbage + no_sewage + gt1hh + remit + drinktap + rental | 2921.97 | 0.02 |
| x ~ 1 + postsec + prim + afro + under15 + headhh_age + housecond + no_garbage + no_piped + no_sewage + gt1hh + remit + drinktap + rental | 2922.00 | 0.02 |
| x ~ 1 + postsec + prim + afro + under15 + under1 +<br>headhh_age + housecond + no_garbage + no_sewage + gt1hh +<br>remit + drinktap + rental | 2922.05 | 0.02 |
| x ~ 1 + postsec + prim + afro + under15 + headhh_age +<br>housecond + no_garbage + no_sewage + gt1hh + remit +<br>drinktap | 2922.05 | 0.02 |
| x ~ 1 + postsec + prim + afro + under15 + under1 +<br>headhh_age + housecond + no_garbage + gt1hh + remit | 2922.09 | 0.02 |
| x ~ 1 + postsec + sec + prim + afro + under15 + headhh_age +<br>housecond + no_garbage + gt1hh + remit | 2922.11 | 0.02 |
| x ~ 1 + postsec + prim + afro + under15 + under1 + headhh_age<br>+ housecond + no_garbage + no_sewage + gt1hh + remit | 2922.13 | 0.02 |
| x ~ 1 + postsec + sec + prim + afro + under15 + headhh_age +<br>housecond + no_garbage + no_sewage + gt1hh + remit | 2922.24 | 0.02 |
| x ~ 1 + postsec + prim + afro + under15 + headhh_age +<br>housecond + no_garbage + no_piped + gt1hh + remit | 2922.26 | 0.02 |
| x ~ 1 + postsec + prim + afro + under15 + headhh_age +<br>pplperhh + housecond + no_garbage + no_sewage + gt1hh +<br>remit + drinktap + rental | 2922.44 | 0.01 |
| x ~ 1 + postsec + prim + afro + under15 + headhh_age +<br>housecond + no_garbage + no_pave + gt1hh + remit | 2922.47 | 0.01 |
| x ~ 1 + postsec + prim + afro + under15 + headhh_age +<br>housecond + no_garbage + no_sewage + gt1hh | 2922.47 | 0.01 |
| x ~ 1 + postsec + prim + afro + under15 + headhh_age +<br>fourpplbedrm + housecond + no_garbage + gt1hh + remit +<br>drinktap + rental | 2922.53 | 0.01 |
| x ~ 1 + postsec + prim + afro + under15 + headhh_age +<br>womhead + housecond + no_garbage + no_sewage + gt1hh +<br>remit + drinktap | 2922.56 | 0.01 |
| x ~ 1 + postsec + prim + afro + under15 + headhh_age +<br>pplperhh + housecond + no_garbage + gt1hh + remit | 2922.63 | 0.01 |

**Supplemental Data File 1.** 2012 dengue case data aggregated to census block level, Guayaquil, Ecuador, provided as a polygon shapefile created in ArcGIS version 10.3.1 (*ArcGIS 10.3.1*, 2016). Census block outlines were digitized by INAMHI (National Institute of Meterology and Hydrology) authors during the course of this project, as reported in the methods.

## References

Alava, A., Mosquera, C., Vargas, W., & Real, J. (2005). Dengue en el Ecuador 1989-2002.Revista Ecuatoriana de Higiene y Medicina Tropical, 42, 11–34.

Almeida, A. S. de, Medronho, R. de A., & Valencia, L. I. O. (2009). Spatial analysis of dengue and the socioeconomic context of the city of Rio de Janeiro (Southeastern Brazil). Revista De Saúde Pública, 43(4), 666–673.

Anselin, L. (2010). Local Indicators of Spatial Association-LISA. Geographical Analysis, 27(2), 93–115. https://doi.org/10.1111/j.1538-4632.1995.tb00338.x

ArcGIS 10.3.1. (2016). (Version 10.3.1). Redlands, CA: Environmental Systems Research

Institute (ESRI). Banaitiene, N. (Ed.). (2012). Risk Management - Current Issues and Challenges. InTech. Retrieved from http://www.intechopen.com/books/risk-management-current-issues-andchallenges

Bertoli, S., Fernandez-Huertas Moraga, J., & Ortega, F. (2011). Immigration Policies and the Ecuadorian Exodus. The World Bank Economic Review, 25(1), 57–76. https://doi.org/10.1093/wber/lhr004

Bertoli, Simone, & Marchetta, F. (2014). Migration, Remittances and Poverty in Ecuador. The Journal of Development Studies, 50(8), 1067–1089. https://doi.org/10.1080/00220388.2014.919382

Bradley, J., Rehman, A. M., Schwabe, C., Vargas, D., Monti, F., Ela, C., … Kleinschmidt, I. (2013). Reduced Prevalence of Malaria Infection in Children Living in Houses with Window Screening or Closed Eaves on Bioko Island, Equatorial Guinea. PLoS ONE, 8(11), e80626. https://doi.org/10.1371/journal.pone.0080626

Burnham, K. P., Anderson, D. R., & Burnham, K. P. (2002). Model selection and multimodel inference: a practical information-theoretic approach (2nd ed). New York: Springer.

Calcagno, V., & Mazancourt, C. de. (2010). glmulti: An R Package for Easy Automated Model Selection with (Generalized) Linear Models. Journal of Statistical Software, 34(12). https://doi.org/10.18637/jss.v034.i12

Castillo, K. C., Körbl, B., Stewart, A., Gonzalez, J. F., & Ponce, F. (2011). Application of spatial analysis to the examination of dengue fever in Guayaquil, Ecuador. Procedia Environmental Sciences, 7, 188–193. https://doi.org/10.1016/j.proenv.2011.07.033

Castillo, Katty C. (2011). Zeit-Raumanalyse des Einflusses von Umwelt und sozialen Faktoreauf den Ausbruch von Denguefieber im Zeitraum 2005-2009 in Guayaquil-Ecuador (Dissertation). Heinrich-Heine-Universität Düsseldorf.

Cazelles, B., Chavez, M., McMichael, A. J., & Hales, S. (2005). Nonstationary Influence of El Niño on the Synchronous Dengue Epidemics in Thailand. PLoS Medicine, 2(4), e106. https://doi.org/10.1371/journal.pmed.0020106

Cummings, D. A. T., Irizarry, R. A., Huang, N. E., Endy, T. P., Nisalak, A., Ungchusak, K., & Burke, D. S. (2004). Travelling waves in the occurrence of dengue haemorrhagic fever in Thailand. Nature, 427(6972), 344–347. https://doi.org/10.1038/nature02225

de Mattos Almeida, M. C., Caiaffa, W. T., Assunção, R. M., & Proietti, F. A. (2007). Spatial Vulnerability to Dengue in a Brazilian Urban Area During a 7-Year Surveillance. Journal of Urban Health, 84(3), 334–345. https://doi.org/10.1007/s11524-006-9154-2

Dick, O. B., Martín, J. L. S., Montoya, R. H., Diego, J. del, Zambrano, B., & Dayan, G. H. (2012). The History of Dengue Outbreaks in the Americas. The American Journal of Tropical Medicine and Hygiene, 87(4), 584–593. https://doi.org/10.4269/ajtmh.2012.11-0770

Endy, T. P., Nisalak, A., Chunsuttiwat, S., Libraty, D. H., Green, S., Rothman, A. L., … Ennis, F. A. (2002). Spatial and Temporal Circulation of Dengue Virus Serotypes: A Prospective Study of Primary School Children in Kamphaeng Phet, Thailand. American Journal of Epidemiology, 156(1), 52–59. https://doi.org/10.1093/aje/kwf006

Galli, B., & Chiaravalloti Neto, F. (2008). Temporal-spatial risk model to identify areas at highrisk for occurrence of dengue fever. Revista De Saúde Pública, 42(4), 656–663.

Global Administrative Areas (2015). (2015). In GADM database of Global Administrative Areas, version 2.8. Retrieved from www.gadm.org.

Hartter, J., Ryan, S. J., MacKenzie, C. A., Parker, J. N., & Strasser, C. A. (2013). Spatially Explicit Data: Stewardship and Ethical Challenges in Science. PLoS Biology, 11(9), e1001634. https://doi.org/10.1371/journal.pbio.1001634

Hiscox, A., Khammanithong, P., Kaul, S., Sananikhom, P., Luthi, R., Hill, N., … Lindsay, S. W. (2013). Risk Factors for Mosquito House Entry in the Lao PDR. PLoS ONE, 8(5), e62769.https://doi.org/10.1371/journal.pone.0062769

Hu, W., Clements, A., Williams, G., & Tong, S. (2011). Spatial analysis of notified dengue fever infections. Epidemiology and Infection, 139(03), 391–399. https://doi.org/10.1017/S0950268810000713

Huang, B., Banzon, V. F., Freeman, E., Lawrimore, J., Liu, W., Peterson, T. C., … Zhang, H.-M.(2015). Extended Reconstructed Sea Surface Temperature Version 4 (ERSST.v4). Part I:Upgrades and Intercomparisons. Journal of Climate, 28(3), 911–930.https://doi.org/10.1175/JCLI-D-14-00006.1

INEC. (2010). Censo de Población y Vivienda. Instituto Nacional de Estadística y Censos.

Jokisch, B., & Pribilsky, J. (2002). The Panic to Leave: Economic Crisis and the “New Emigration” from Ecuador. International Migration, 40(4), 75–102. https://doi.org/10.1111/1468-2435.00206

Kanamitsu, M., Ebisuzaki, W., Woollen, J., Yang, S.-K., Hnilo, J. J., Fiorino, M., & Potter, G. L.(2002). NCEP–DOE AMIP-II Reanalysis (R-2). Bulletin of the American Meteorological Society, 83(11), 1631–1643. https://doi.org/10.1175/BAMS-83-11-1631

Kuhn, Katrin, Campbell-Lendrum, Diarmid, Haines, Andy, & Cox, Jonathan. (2005). Using climate to predict infectious disease epidemics. World Health Organization. Retrieved from http://www.who.int/globalchange/publications/infectdiseases.pdf?ua=1

Ministerio de Salud Pública. (2013). Boletín epidemiológico de la situación del Dengue en el Ecuador., (No. 46). Retrieved from http://www.salud.gob.ec/boletin-epidemiologico-de-lasituacion-del-dengue-en-el-ecuador-no-46-07-de-enero-de-2013/

Mondini, A., Bronzoni, R. V. de M., Nunes, S. H. P., Chiaravalloti Neto, F., Massad, E., Alonso, W. J., … Nogueira, M. L. (2009). Spatio-Temporal Tracking and Phylodynamics of an Urban Dengue 3 Outbreak in São Paulo, Brazil. PLoS Neglected Tropical Diseases, 3(5), e448. https://doi.org/10.1371/journal.pntd.0000448

Mulligan, K., Dixon, J., Joanna Sinn, C.-L., & Elliott, S. J. (2015). Is dengue a disease of poverty? A systematic review. Pathogens and Global Health, 109(1), 10–18. https://doi.org/10.1179/2047773214Y.0000000168

Muñoz, Á. G., Thomson, M. C., Goddard, L., & Aldighieri, S. (2016). Analyzing climate variations at multiple timescales can guide Zika virus response measures. GigaScience, 5(1). https://doi.org/10.1186/s13742-016-0146-1

Muñoz, Á.G., Stewart-Ibarra, A.M., & Ruiz-Carrascal, D. (2013). Desarrollo de Modelos de Pronóstico Experimental: Análisis Socio Ecológico de Riesgo a Dengue y Análisis Estadístico de Patrones Climáticos, Entomológicos y Epidemiológicos en Modelos de Dengue. Technical Report, CLIDEN Project. INAMHI-SENESCYT, Quito, Ecuador.

Oliveira, M. A. de, Ribeiro, H., & Castillo-Salgado, C. (2013). Geospatial analysis applied to epidemiological studies of dengue: a systematic review. Revista Brasileira De Epidemiologia = Brazilian Journal of Epidemiology, 16(4), 907–917.

Padmanabha, H., Durham, D., Correa, F., Diuk-Wasser, M., & Galvani, A. (2012). The Interactive Roles of Aedes aegypti Super-Production and Human Density in Dengue Transmission. PLoS Neglected Tropical Diseases, 6(8), e1799. https://doi.org/10.1371/journal.pntd.0001799

Pan American Health Organization, & World Health Organization. (2016). Zika suspected and confirmed cases reported by countries and territories in the Americas. Cumulative cases, 2015 - 2016. PAN/WHO, Washington, D.C. Retrieved from http://www.paho.org/hq/index.php?option=com_content&view=article&id=12390&Itemid=42090&lang=en

Real, J., & Mosquera, C. (2012). Detección del Virus Dengue en el Ecuador. Una Vision Epidemiologica. Período 1988 - 2012. Instituto Nacional de Higiene y Medicina Tropical.

Reiner, R. C., Stoddard, S. T., Forshey, B. M., King, A. A., Ellis, A. M., Lloyd, A. L., … Scott, T. W. (2014). Time-varying, serotype-specific force of infection of dengue virus. Proceedings of the National Academy of Sciences, 111(26), E2694–E2702. https://doi.org/10.1073/pnas.1314933111

Rodrigues, M., Marques, G., Serpa, L., Arduino, M., Voltolini, J., Barbosa, G., … de Lima, V. (2015). Density of Aedes aegypti and Aedes albopictus and its association with number of residents and meteorological variables in the home environment of dengue endemic area, São Paulo, Brazil. Parasites & Vectors, 8(1), 115. https://doi.org/10.1186/s13071-015-0703-y

San Martín, J. L., Brathwaite, O., Zambrano, B., Solórzano, J. O., Bouckenooghe, A., Dayan, G.H., & Guzmán, M. G. (2010). The Epidemiology of Dengue in the Americas Over the Last Three Decades: A Worrisome Reality. The American Journal of Tropical Medicine and Hygiene, 82(1), 128–135. https://doi.org/10.4269/ajtmh.2010.09-0346

Schmidt, W.-P., Suzuki, M., Dinh Thiem, V., White, R. G., Tsuzuki, A., Yoshida, L.-M., … Ariyoshi, K. (2011). Population Density, Water Supply, and the Risk of Dengue Fever in Vietnam: Cohort Study and Spatial Analysis. PLoS Medicine, 8(8), e1001082. https://doi.org/10.1371/journal.pmed.1001082

Stewart Ibarra, A. M., Luzadis, V. A., Borbor Cordova, M. J., Silva, M., Ordoñez, T., Beltrán Ayala, E., & Ryan, S. J. (2014). A social-ecological analysis of community perceptions of dengue fever and Aedes aegypti in Machala, Ecuador. BMC Public Health, 14(1). https://doi.org/10.1186/1471-2458-14-1135

Stewart Ibarra, A. M., Ryan, S. J., Beltrán, E., Mejía, R., Silva, M., & Muñoz, Á. (2013). Dengue Vector Dynamics (Aedes aegypti) Influenced by Climate and Social Factors in Ecuador: Implications for Targeted Control. PLoS ONE, 8(11), e78263. https://doi.org/10.1371/journal.pone.0078263

Stewart-Ibarra, A. M., Muñoz, Á. G., Ryan, S. J., Ayala, E. B., Borbor-Cordova, M. J., Finkelstein, J. L., … Rivero, K. (2014). Spatiotemporal clustering, climate periodicity, and social-ecological risk factors for dengue during an outbreak in Machala, Ecuador, in 2010. BMC Infectious Diseases, 14(1). https://doi.org/10.1186/s12879-014-0610-4

Stoddard, S. T., Morrison, A. C., Vazquez-Prokopec, G. M., Paz Soldan, V., Kochel, T. J., Kitron, U., … Scott, T. W. (2009). The Role of Human Movement in the Transmission of Vector-Borne Pathogens. PLoS Neglected Tropical Diseases, 3(7), e481. https://doi.org/10.1371/journal.pntd.0000481

Teixeira, M. da G., Barreto, M. L., Costa, M. da C. N., Ferreira, L. D. A., Vasconcelos, P. F. C., & Cairncross, S. (2002). Dynamics of dengue virus circulation: a silent epidemic in a complex urban area. Tropical Medicine & International Health: TM & IH, 7(9), 757–762.

Thomson, M. C., García Herrera, R., & Beniston, M. (2008). Seasonal forecasts, climatic change and human health: health and climate. Dordrecht; London: Springer. Retrieved from http://site.ebrary.com/id/11029199

US Military, Department of Defense (all branches). (1992). Digital Chart of the World (DCW). USA.

Van Benthem, B. H. B., Khantikul, N., Panart, K., Kessels, P. J., Somboon, P., & Oskam, L. (2002). Knowledge and use of prevention measures related to dengue in northern Thailand. Tropical Medicine & International Health: TM & IH, 7(11), 993–1000.

Van Benthem, Birgit H. B., Vanwambeke, S. O., Khantikul, N., Burghoorn-Maas, C., Panart, K., Oskam, L., … Somboon, P. (2005). Spatial patterns of and risk factors for seropositivity for dengue infection. The American Journal of Tropical Medicine and Hygiene, 72(2), 201–208.

Vazquez-Prokopec, G. M., Kitron, U., Montgomery, B., Horne, P., & Ritchie, S. A. (2010). Quantifying the Spatial Dimension of Dengue Virus Epidemic Spread within a Tropical Urban Environment. PLoS Neglected Tropical Diseases, 4(12), e920. https://doi.org/10.1371/journal.pntd.0000920

Vazquez-Prokopec, G. M., Stoddard, S. T., Paz-Soldan, V., Morrison, A. C., Elder, J. P., Kochel, T. J., … Kitron, U. (2009). Usefulness of commercially available GPS data-loggers for tracking human movement and exposure to dengue virus. International Journal of Health Geographics, 8(1), 68. https://doi.org/10.1186/1476-072X-8-68

Who | Global Strategy for dengue prevention and control, 2012–2020. (n.d.). Retrieved May 5, 2014, from http://www.who.int/denguecontrol/9789241504034/en/

Yoon, I.-K., Getis, A., Aldstadt, J., Rothman, A. L., Tannitisupawong, D., Koenraadt, C. J. M., … Scott, T. W. (2012). Fine Scale Spatiotemporal Clustering of Dengue Virus Transmission in Children and Aedes aegypti in Rural Thai Villages. PLoS Negl Trop Dis, 6(7), e1730.https://doi.org/10.1371/journal.pntd.0001730

Zambrano, H., Waggoner, J. J., Almeida, C., Rivera, L., Benjamin, J. Q., & Pinsky, B. A. (2016). Zika Virus and Chikungunya Virus CoInfections: A Series of Three Cases from a Single Center in Ecuador. American Journal of Tropical Medicine and Hygiene. https://doi.org/10.4269/ajtmh.16-0323

